# Discovery of D2469079A, A novel selective Toll-Like Receptor 7/8 Antagonist with Brain Penetrance

**DOI:** 10.1101/2025.04.12.648517

**Authors:** Xiaobing Deng, Xun Zhang, Wenjia Tian, Donghuai Xiao, Zhichao Du, Yanhong Wang, Shou Cao, Bo Li, Yan Zhao, Huiling Chen, Yaning Su, Yisui Zhou

**Affiliations:** Beijing double-crane runchuang technology Co. Ltd., Beijing, China

**Keywords:** Toll-like receptor7, toll-like receptor 8, lupus, innate immunity, brain permeability

## Abstract

Aberrant activation of endosomal Toll-like receptors (TLRs) 7 and 8, which recognize single-stranded RNA, is strongly implicated in the pathogenesis of autoimmune diseases such as systemic lupus erythematosus (SLE). And the frequent neuropsychiatric involvement in SLE highlights the need for brain-penetrant therapies. Here, we report the identification and characterization of D2469079A, a novel, potent dual TLR7/TLR8 antagonist (low nM IC_50_ in humanized HEK-Blue assays) selective against hTLR9. D2469079A demonstrates favorable *in vitro* properties, including adequate solubility and high microsomal stability. It exhibited desirable pharmacokinetic profiles across multiple preclinical species (mice, rats, dogs) and achieved significant cerebrospinal fluid (CSF) exposure, confirming its high permeability and brain penetration. Consistent *in vivo* efficacy was observed in both acute and chronic stimulation models. Importantly, D2469079A displayed a favorable safety profile, lacking significant liabilities related to CYP450 inhibition/induction, hERG interaction, or toxicity in a 14-day rat study. These findings position D2469079A as a promising drug candidate with excellent drug-like characteristics for the treatment of TLR7/8-driven autoimmune diseases, potentially including neuropsychiatric manifestations. We are actively seeking partners for collaborative clinical development.

---

Toll-like receptors (TLRs) are a class of molecular pattern recognition receptors widely distributed across various tissues. They monitor and recognize different pathogen-associated molecular patterns (PAMPs) and damage-associated molecular patterns (DAMPs), playing a crucial role in both innate and adaptive immunity. (*1*) Endosomal TLRs, including TLR3, TLR7/8, and TLR9, can respectively recognize double-stranded RNA (dsRNA), GU-rich single-stranded RNA (ssRNA), and unmethylated single-stranded DNA (ssDNA). The aberrant activation of these nucleic acid-sensing endosomal TLRs is considered a potential driving factor in autoimmune diseases such as systemic lupus erythematosus (SLE), Sjögren’s syndrome (SjS), systemic sclerosis (SSc), and derma tomyositis (DM)(*2-4*). Research has identified a human SLE case associated with a gain-of-function mutation in TLR7, providing the first validation of a causal relationship between TLR7 and SLE in humans(*5*), and an early-onset SLE caused by a somatic TLR7 gain-of-function mutation further confirmed the central role of TLR7 in disease pathogenesis(*6*). Preliminary clinical results of BMS986256 and M5049 demonstrate the therapeutic potential of targeting TLR7 for CLE/SLE(*7, 8*). The nervous system is one of the major organs affected in patients with SLE(*9*), the true prevalence of neuropsychiatric SLE is unknown, but published estimates suggest that it affects between 12% and 95% of patients with SLE(*9-13*). Here we report a highly permeable, selective, and brain-penetrant TLR7/8 antagonist for further clinical development.

## Profiling Data of Compounds

**Table 1.**
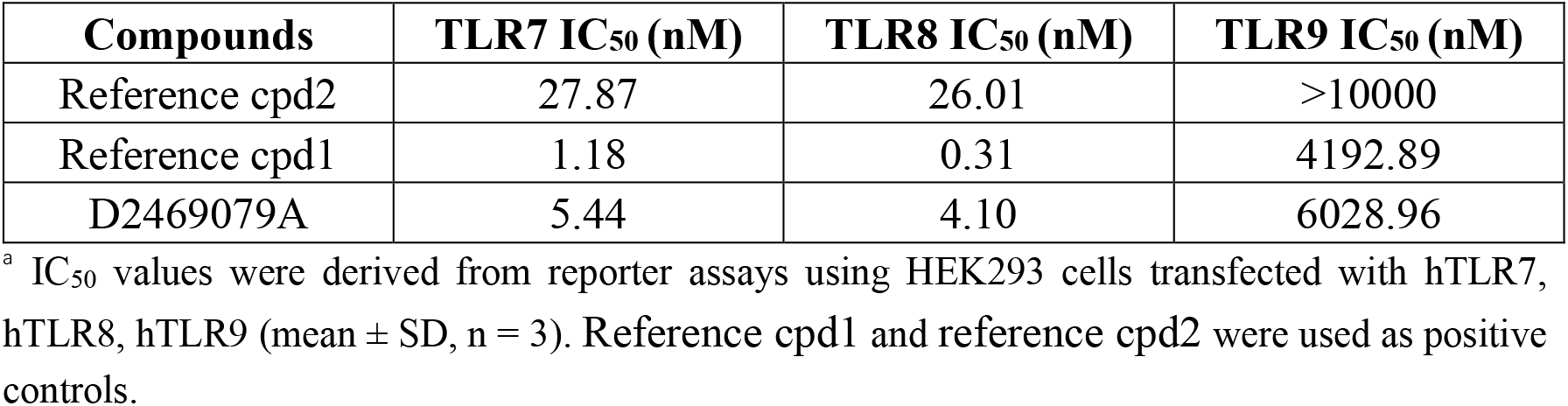
Effects on TLR7, TLR8 and TLR9 Antagonism^a^.

### Solubility

The compound D2469079A showed good solubility(Table 2)

**Table 2.**
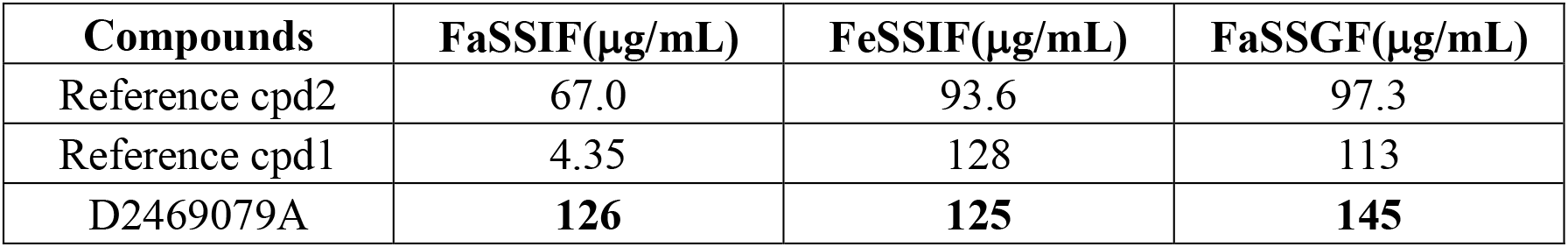
Solubility in different solvents.

### PAMPA

**Table 3.**
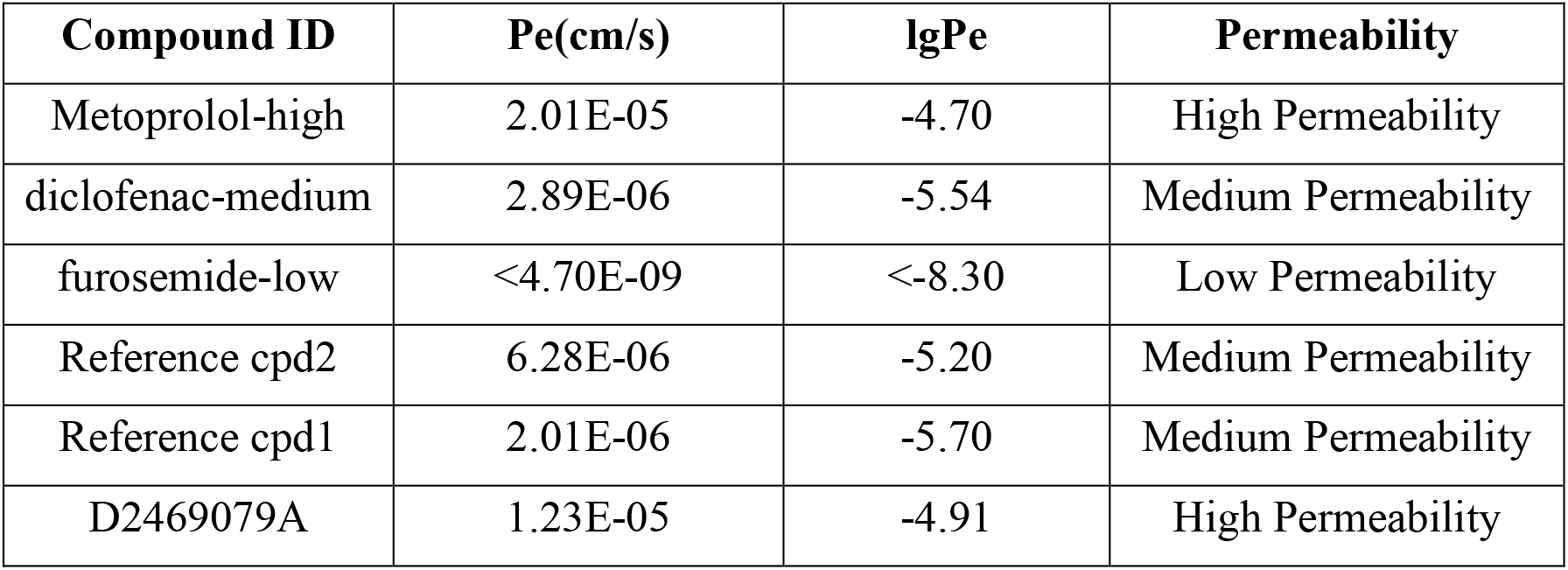
PAMPA permeability of the Antagonists.

**Table 4.**
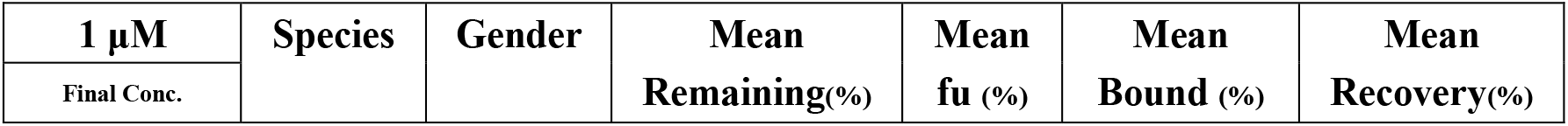
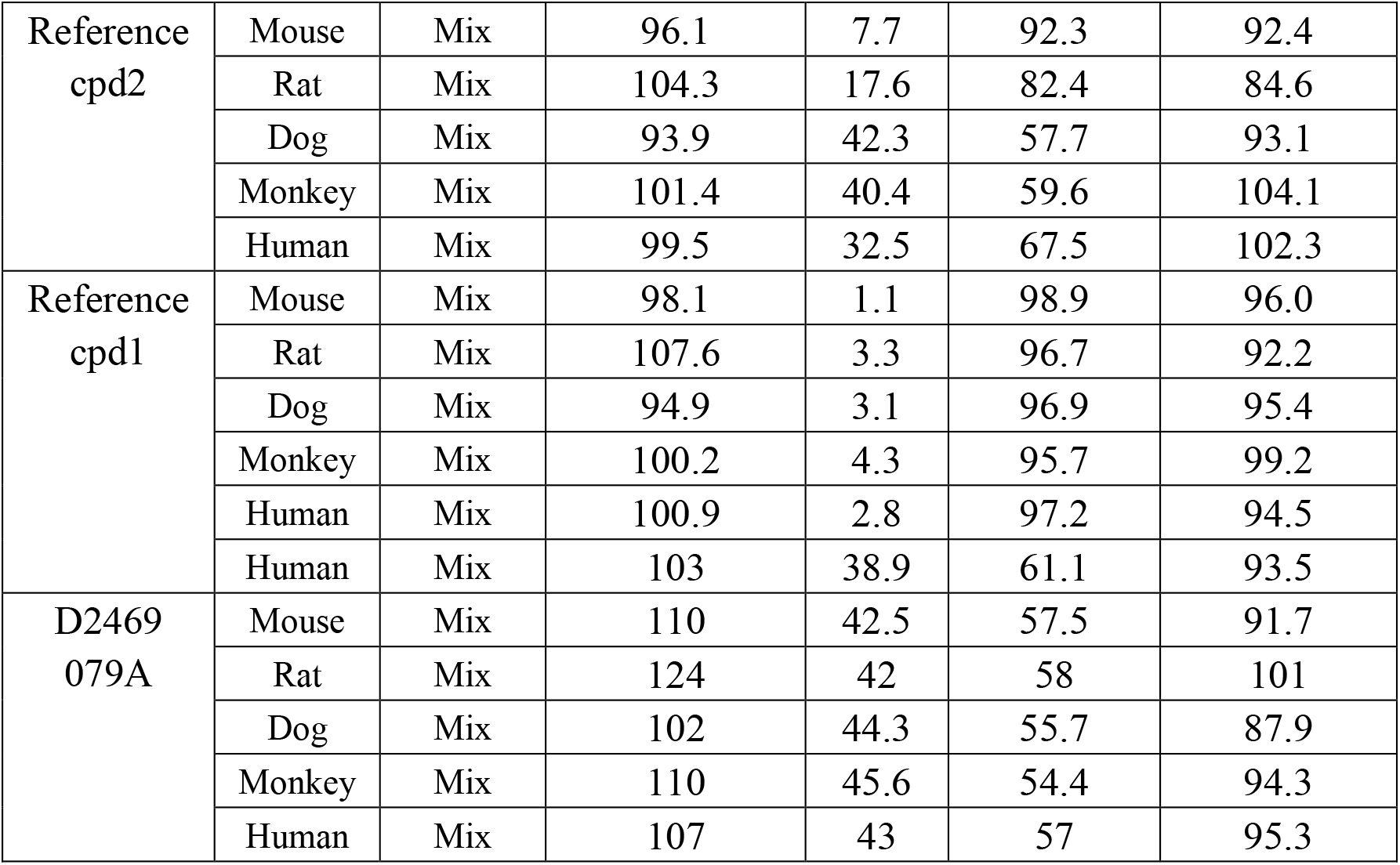
Plasma protein binding.

|  |  |  |  |  |  |  |
| --- | --- | --- | --- | --- | --- | --- |
| Reference<br>cpd2 | Mouse | Mix | 96.1 | 7.7 | 92.3 | 92.4 |
|  | Rat | Mix | 104.3 | 17.6 | 82.4 | 84.6 |
|  | Dog | Mix | 93.9 | 42.3 | 57.7 | 93.1 |
|  | Monkey | Mix | 101.4 | 40.4 | 59.6 | 104.1 |
|  | Human | Mix | 99.5 | 32.5 | 67.5 | 102.3 |
| Reference<br>cpd1 | Mouse | Mix | 98.1 | 1.1 | 98.9 | 96.0 |
|  | Rat | Mix | 107.6 | 3.3 | 96.7 | 92.2 |
|  | Dog | Mix | 94.9 | 3.1 | 96.9 | 95.4 |
|  | Monkey | Mix | 100.2 | 4.3 | 95.7 | 99.2 |
|  | Human | Mix | 100.9 | 2.8 | 97.2 | 94.5 |
|  | Human | Mix | 103 | 38.9 | 61.1 | 93.5 |
| D2469<br>079A | Mouse | Mix | 110 | 42.5 | 57.5 | 91.7 |
|  | Rat | Mix | 124 | 42 | 58 | 101 |
|  | Dog | Mix | 102 | 44.3 | 55.7 | 87.9 |
|  | Monkey | Mix | 110 | 45.6 | 54.4 | 94.3 |
|  | Human | Mix | 107 | 43 | 57 | 95.3 |

**Table 5.** *in vitro* Pharmacokinetic Profile of the selected compounds.

| **1 μM**<br>Final Conc | **Species** | **Gender** | **T60min**<br>**Remaining(%)** | **t<sub>1/2</sub> (min)** | **Clint**<br>(uL/min/mg) | **ER** |
| --- | --- | --- | --- | --- | --- | --- |
| Reference<br>cpd2 | Mouse | M | 100.2 | > 186 | < 7.44 | < 0.25 |
|  | Rat | M | 106.23 | > 186 | < 7.44 | < 0.20 |
|  | Dog | M | 103.01 | > 186 | < 7.44 | < 0.30 |
|  | Monkey | M | 98.57 | > 186 | < 7.44 | < 0.17 |
|  | Human | Mix | 102.01 | > 186 | < 7.44 | < 0.27 |
| Reference<br>cpd1 | Mouse | M | 85.69 | > 186 | < 7.44 | < 0.25 |
|  | Rat | M | 64.09 | 98.0 | 14.1 | 0.3 |
|  | Dog | M | 71.74 | 126.8 | 10.9 | 0.4 |
|  | Monkey | M | 68.38 | 112.7 | 12.3 | 0.4 |
|  | Human | Mix | 87.67 | > 186 | < 7.44 | < 0.27 |
| D2469<br>079A | Mouse | M | 90.32 | > 186 | < 7.44 | < 0.25 |
|  | Rat | M | 80 | > 186 | < 7.44 | < 0.20 |
|  | Dog | M | 84.27 | > 186 | < 7.44 | < 0.30 |
|  | Monkey | M | 66.84 | 106 | 13.1 | 0.26 |
|  | Human | Mix | 86.95 | > 186 | < 7.44 | < 0.27 |

**Table 6.** *in vivo* Blood pharmacokinetic profile of D2469079A.

| **Parameter** | **Mice**<br>**10 mg/kg P.O.** | **Rats**<br>**25mg/kg P.O.** | **Dogs**<br>**1 mg/kg P.O.** |
| --- | --- | --- | --- |
| T<sub>1/2</sub> (h) | - | 2.61 | 6.26 |
| T<sub>max</sub>(h) | 1.67 | 1.00 | 3.00 |
| C<sub>max</sub> (ng/ml) | 818 | 930 | 101 |
| AUC last (ng·h/ml) | 3046 | 4362 | 1046 |
| F% | 59.4 | 63.7 | - |

**Table 7.** The CSF drug concentration of D2469079A.

| **D2469079A**<br>**P.O.**<br>**10 mg/kg** | **Time after dosing** | **CSF drug (ng/mL)** |
| --- | --- | --- |
|  | 0.5h | 19.11 |
|  | 2h | 21.15 |
|  | 6h | 7.04 |

**Table 8.** CYP450 Inhibition profile of D2469079A.

| **CYP**<br>**P450 isoforms** | **%Inhibition (10μM Final Conc. )** |  |  |  |  |  |  |
| --- | --- | --- | --- | --- | --- | --- | --- |
|  | **CYP1A2** | **CYP2B6** | **CYP2C8** | **CYP2C9** | **CYP2C19** | **CYP2D6** | **CYP3A_T** |
| PC Inhibitors | 73.8 | 91.4 | 86.7 | 93.7 | 93.9 | 93.7 | 97.3 |
| Reference cpd2 | 7.47 | 1.40 | 15.92 | 12.46 | 7.69 | 61.28 | 2.38 |
| Reference cpd1 | 10.90 | 0.00 | 7.96 | 25.24 | 9.16 | 15.13 | 6.35 |
| D2469079A | 0.0 | 10.5 | 12.4 | 12.8 | 0.0 | 1.4 | 4.1 |

**Figure 1:**
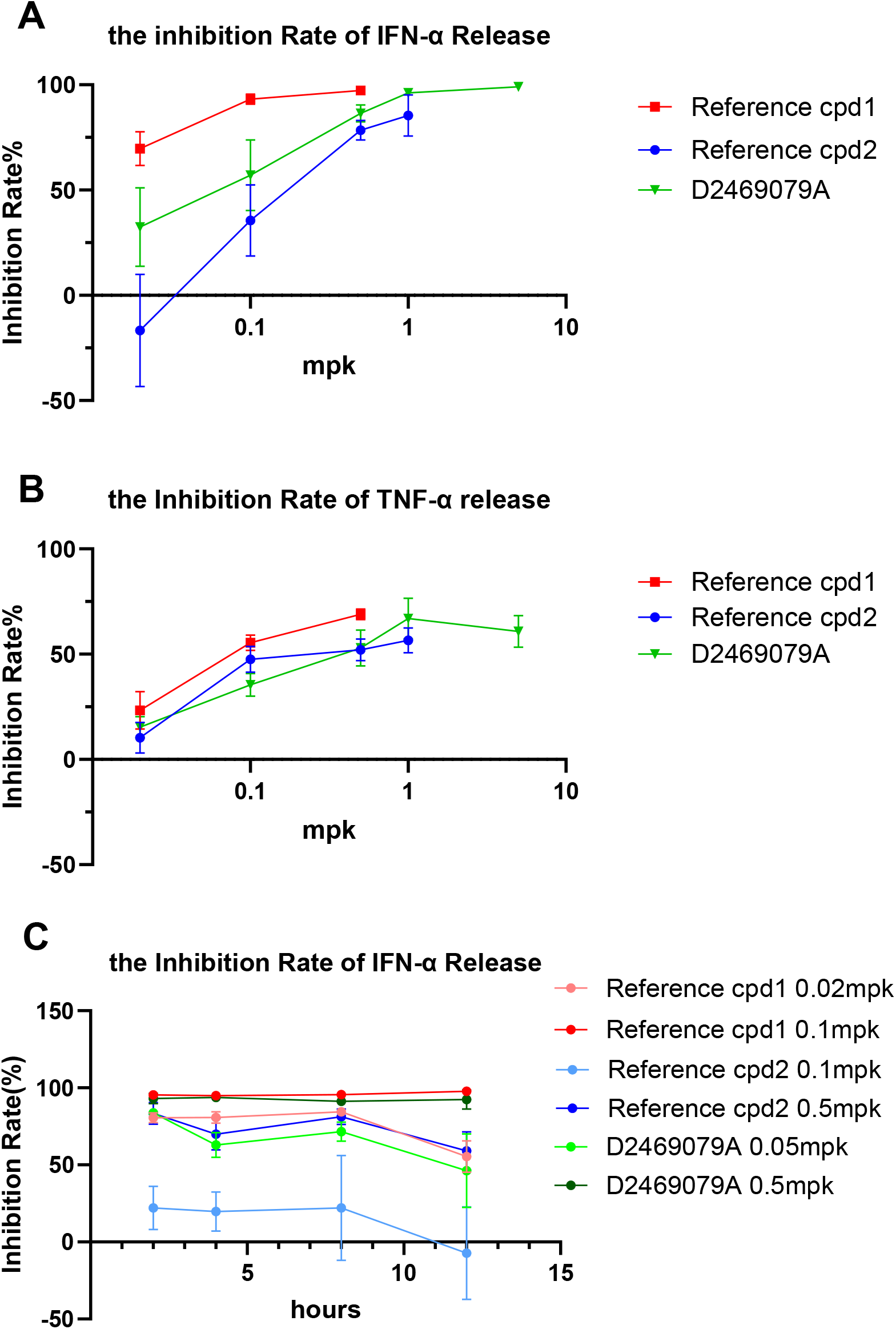
In Vivo TLR7 Antagonism Efficacy. (A) Dose-dependent antagonistic response to acute TLR7 agonist stimulation. (B).Dose-dependent antagonistic response to acute TLR7 agonist stimulation.(C). Dose and time dependence of the antagonistic response to acute TLR7 agonist stimulation. Data are shown as mean ± SEM, n = 4.

**Figure 2:**
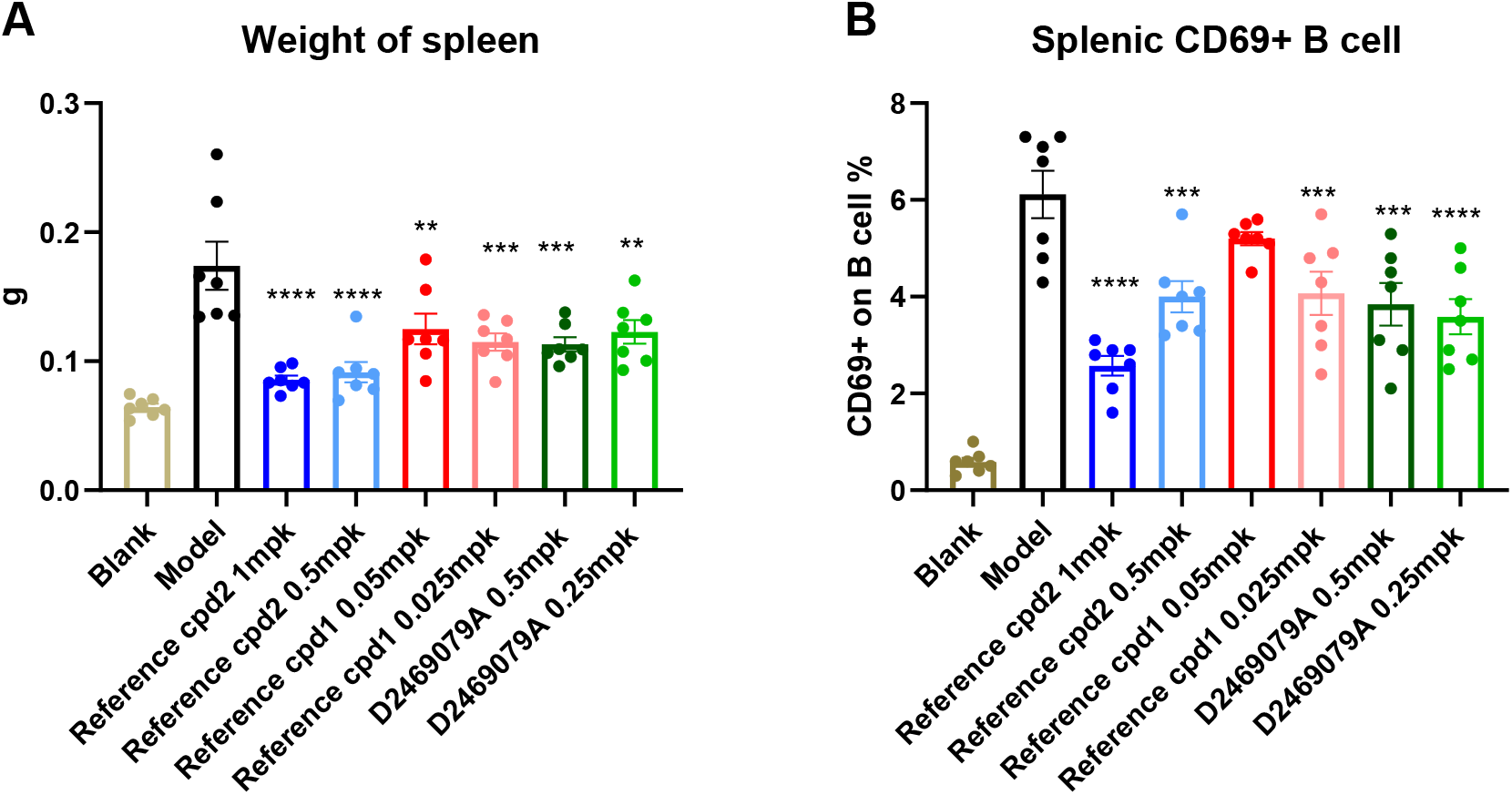
In Vivo TLR7 Antagonism Efficacy of 14-day. ( A).Antagonistic response to 14-day chronic TLR7 agonist stimulation. (B) Antagonistic response to 14-day chronic TLR7 agonist stimulation. Data are shown as mean ± SEM, n = 7.

## Conclusion

Herein, we report the identification of D2469079A, a potent and selective dual TLR7/TLR8 inhibitor with no activity against TLR9. This compound possesses favorable *in vitro* properties, including potent inhibition, adequate solubility, and high hepatic microsomal stability. D2469079A exhibited desirable pharmacokinetic profiles across mice, rats, and dogs, and achieved significant CSF exposure owing to its high permeability. Consistent *in vivo* efficacy was demonstrated in both acute and chronic stimulation models. Importantly, safety evaluations revealed no significant liabilities, including no risks associated with CYP450 inhibition or induction, or hERG channel interaction, and no notable toxicity in a 14-day rat study. Taken together, D2469079A displays excellent drug-like characteristics, positioning it as a promising drug candidate for therapies targeting endosomal TLR7/8. **We are seeking potential partners for collaborative clinical development. Interested parties are encouraged to contact us at:**.

## EXPERIMENTAL SECTION

### Reagents and Materials

Reference cpd1 was example No.15 from BMS patent WO2018005586A1, and reference cpd2 was M5049 from Merck publication(*14*).

HPLC grade methanol and acetonitrile were purchased from Thermo Fisher (Waltham, MA, USA). DMSO, acetic acid, ammonium formate, diclofenac sodium, warfarin, verapamil hydrochloride, α-naphthoflavone, ticlopidine HCl, montelukast sodium, sulfaphenazole, (S)-(+)-N-3-benzylnirvanol, quinidine, and ketoconazole were purchased from Sigma-Aldrich (St. Louis, MO, USA). FaSSIF, FeSSIF, and FaSSGF powders were obtained from Biorelevant (London, UK).

Human liver microsomes and EDTA-K_2_ plasma were purchased from IPHASE (Beijing, China). NADPH was obtained from Solarbio (Beijing, China). MultiScreen® HTS HV 0.45 µm filter plates, PTFE acceptor plates and MultiScreen-IP PAMPA assay plates were purchased from Millipore (Burlington, MA, USA).

Lecithin was purchased from Yuanye (Shanghai, China), and dodecane was obtained from Alfa Aesar (Haverhill, MA, USA). The dialysis membranes (12-14 kDa) and HTD 96b dialysis device were purchased from HTDialysis (Gales Ferry, CT, USA). Phosphate buffer (100 mM, pH 7.4) and 50% acetonitrile water was prepared in-house. The LC-MS/MS system (Sciex Triple Quad 6500) was used for sample analysis. Centrifugation was performed using an Eppendorf 5804R centrifuge (Hamburg, Germany).

MS grade methanol and acetonitrile were purchased from Thermo Fisher (Waltham, MA, USA). DMSO, EDTA-K2 plasma were purchased from IPHASE (Beijing, China). 50% acetonitrile water was prepared in-house.The LC-MS/MS system (Sciex Triple Quad 6500) was used for sample analysis. Centrifugation was performed using an Eppendorf 5910R centrifuge (Hamburg, Germany).

### Reporter Assay to Measure, TLR7, TLR8 and TLR9 Antagonism

HEK293 cells were stably transfected with either TLR7, TLR8 or TLR9 and an NF-κB-luciferase reporter gene. For compound testing, cells were seeded in 384-well (Corning), and after an overnight incubation at 37°C and 5% CO_2_. HEK-Blue cells (invivogen)were treated with increasing concentrations of test compounds (D2469079A, reference cpd1, reference cpd1) or vehicle control (0.1% v/v DMSO) and stimulated with R848 or ODN2006 for 24 h (250ng/mL R848 for TLR7; 2μg/mL for TLR8; 100ng/mL ODN2006 for TLR9). 2 μL of cell culture supernatant was transferred and mixed with 18 μL/well Quanti-Blue® detection reagent (InvivoGen). After 1-hour incubation at room temperature, luminescence signals were measured using a microplate reader (BMG).

### Kinetic Solubility Assay

Test compounds and diclofenac sodium (positive control) were prepared as 10 mM stock solutions in DMSO. For solubility determination, 30 μL of each stock solution was mixed with 970 μL of FaSSIF, FeSSIF or FaSSGF buffer (pre-adjusted to pH 6.5, 5.0 or 1.6) in duplicate. The mixtures were shaken at 37°C (1200 rpm) for 24 hours, then filtered through 0.45 μm membranes. Filtrates were diluted with 50% acetonitrile water (1:100), vortexed, centrifuged, and analyzed by LC-MS/MS. Standard Sample (3 μM) were prepared for quantification. Solubility was calculated based on analyte-to-internal standard peak area ratios.

### PAMPA Permeability Assay

Stock solutions (1 mM in DMSO) of test compounds and permeability controls (metoprolol, diclofenac, furosemide) were diluted to 10 μM in PBS (1% DMSO). Artificial membranes were prepared fresh by dissolving lecithin in dodecane (18 mg/mL). In the assay, 5 μL membrane solution was applied to filter plates, followed by 300 μL compound solution (donor) and 150 μL PBS/1% DMSO (acceptor). After 16 h incubation at 25°C (60 rpm), samples from both chambers were collected, protein-precipitated with cold acetonitrile (containing dexamethasone IS), and analyzed by LC-MS/MS. Effective permeability (Pe) was calculated using chamber volumes (donor: 0.3 mL; acceptor: 0.15 mL), incubation time (57600 s), and membrane area (0.24 cm^2^).

### Plasma Protein Binding Assay

Test compounds and warfarin (positive control) were prepared as 200 μM working solutions in DMSO, then diluted to 1 μM in EDTA-K2 plasma. Dialysis membranes (12-14 kDa) were pre-treated with water, 20% ethanol, and PBS buffer (pH 7.4). The assay was conducted in duplicate using 120 μL plasma samples dialyzed against equal volumes of PBS at 37°C (100 rpm) for 6 h. After incubation, plasma and buffer samples were protein-precipitated with cold acetonitrile (IS), centrifuged, and analyzed by LC-MS/MS. Percent free drug was calculated from buffer-to-plasma peak area ratios.

### Microsomal Stability Assay

Liver microsomes (20 mg/mL) were diluted to 0.5 mg/mL in phosphate buffer (pH 7.4). Test compounds and verapamil (positive control) were added at 1 μM final concentration. Reactions were initiated with NADPH (1 mM), and then aliquots were taken at 0, 5, 10, 15, 30, and 60 min, quenched with cold acetonitrile (IS), and centrifuged. Supernatants were analyzed by LC-MS/MS to determine remaining parent compound. In vitro half-life (t1/2) and intrinsic clearance (CLint) were calculated from logarithmic plots of concentration versus time.

### CYP450 Inhibition Assay

Human liver microsomes (0.1 mg/mL) were pre-incubated with test compounds and CYP-specific substrates: phenacetin (1A2), bupropion (2B6), amodiaquine (2C8), diclofenac (2C9), (S)-mephenytoin (2C19), dextromethorphan (2D6), and testosterone (3A4). Reactions were initiated with NADPH (1 mM) and stopped at 10 min (30 min for CYP2C19) by acetonitrile precipitation. Metabolite formation was quantified by LC-MS/MS. Inhibition percentage was calculated relative to control reactions without inhibitor. Selective inhibitors (e.g., ketoconazole for CYP3A4) were included as controls.

### Animal Studies

All procedures using animals were performed in accordance with the Guide for the Care and Use of Laboratory Animals and all local and national laws and regulations regarding animal care under protocols approved by the local Institutional Animal Care and Use Committee.

### Pharmacokinetic Study

Mice were treated with 2 mg/kg(iv), 10 mg/kg (p.o.). Rats were treated with 1 mg/kg(iv), 25 mg/kg (p.o.). Dogs were treated with 1 mg/kg(p.o.). For each mouse in all group, blood samples were collected at the time point of 0.25, 0.5, 1, 2, 4, 8 and 24 h post-dose. For each rat in all group, blood samples were collected at the time point of 0.25, 0.5, 1, 2, 4, 8, 12 and 24 h post-dose. For each dog in all group, blood samples were collected at the time point of 0.25, 0.5, 1, 2, 4, 8, 12 and 28 h post-dose Blood samples were sampled into EDTA-coated tubes (Eppendorf) and placed on ice until centrifugation to obtain plasma samples. The plasma samples were stored at -80°C until analysis. The concentration of compound in plasma samples was determined using a LC-MS/MS method. PK parameters were estimated by non-compartmental model using WinNonlin.

### Cerebrospinal Fluid drugs concentration

For D2469079A at each time point, three female Sprague-Dawley (SD) rats were used, with a total of two groups. The rats were treated with a single oral dose of 10 mg/kg (via gavage). For each rat, cerebrospinal fluid (CSF) samples were collected at 0.5, 2, and 6 hours post-dose. The concentrations of the D2469079A in the CSF samples were determined using an LC-MS/MS method.

### In Vivo Inhibition Assay

#### Acute cytokines release *in vivo* assay

Female C57BL/6 mice (8-10 weeks, Charles River Laboratories) were dosed via oral gavage with test compounds (D2469079A, reference cpd1, reference cpd2) formulated in 10% hydroxypropyl-β-cyclodextrin (HP-β-CD) .At predetermined intervals post-administration, mice received intraperitoneal injections (i.p.) of R848 (25 μg) dissolved in 0.9% saline. Two hours post-R848 challenge, the mice were euthanized, and plasma samples were obtained via retro-orbital bleeding into EDTA-coated tubes (BD Biosciences) to inhibit coagulation. Plasma TNF-α and IFN-α concentrations were quantified using enzyme-linked immunosorbent assay (ELISA, Thermo Fisher Scientific) according to manufacturer protocols, with absorbance measured at 450 nm using a SpectraMax M5 microplate reader (Molecular Devices).

#### Chronic TLR7 activation in vivo assay

Female C57BL/6 mice (8-10 weeks, n=5/group) received daily intraperitoneal (i.p.) injections of R848 (2.5 mg/kg) for 14 days. Commencing on day 7, experimental groups were orally administered graded doses of test compounds: D2469079 (0.25 mg/kg and 0.5 mg/kg), reference cpd2 (0.5 mg/kg and 1 mg/kg), and reference cpd1 (0.025 mg/kg and 0.05 mg/kg), while the control group received vehicle (10% HP-β-CD). On day 14, all treatment groups were compared with age-matched naive controls. Primary endpoints included spleen weight measurement and flow cytometric quantification of activated splenic B cells (B220^+^CD69^+^) using a BD LSRFortessa analyzer (BD Biosciences). Splenocytes were stained with APC-anti-B220 (Biolegend) and PE-anti-CD69 (Biolegend) antibodies.

## Data Analysis

For in vitro assays testing compounds (D2469079A, reference cpd1, reference cpd2) against TLR7, TLR8 and TLR9 in HEK-Blue cells, experiments were performed in duplicate or triplicate, and IC_50_ values were calculated using a four-parameter fit of data in Prism (GraphPad, San Diego, CA).

## Notes

### Competing Interest Statement

All authors are employees.

